# Mutation of an L-Type Calcium Channel Gene Leads to a Novel Human Primary Cellular Immunodeficiency

**DOI:** 10.1101/864280

**Authors:** Franz Fenninger, Shawna R. Stanwood, Chieh-Ju Lu, Cheryl G. Pfeifer, Sarah E. Henrickson, Omar Khan, Kaitlin C. O’Boyle, Kelly Maurer, Melanie Ruffner, Ramin S. Herati, Neil D. Romberg, E. John Wherry, Kathleen E. Sullivan, Wilfred A. Jefferies

## Abstract

Human primary immunodeficiencies are inherited diseases that can provide valuable insight into the immune system. Calcium (Ca^2+^) is a vital secondary messenger in T lymphocytes regulating a vast array of important events including maturation, homeostasis, activation, and apoptosis and can enter the cell through CRAC, TRP, and Cav channels. Here we describe three Cav1.4-deficient siblings presenting with X-linked incomplete congenital stationary night blindness as well as an immune phenotype characterized by several recurrent infections. Complete exome sequencing demonstrated that the patients share only a single pathogenic allele; a R625X (p.Arg625Ter) point mutation that leads to a premature stop codon in the *CACNA1F* gene encoding the L-type Ca^2+^ channel Cav1.4. The subjects uniformly exhibited an expansion of central and effector memory T lymphocytes, and evidence of T lymphocytes exhaustion with corresponding upregulation of inhibitory receptors. Moreover, the sustained elevated levels of activation markers on B lymphocytes suggest that they are in a chronic state of activation. Finally, the T lymphocytes from patients and *CACNA1F* knockdown Jurkat T lymphocytes exhibited a reduced Ca^2+^ flux, compared to controls. This is the first example where the mutation of any Cav channel causes a primary immunodeficiency in humans and establishes the physiological importance of Cav channels in the human immune system.

## Introduction

Human primary immunodeficiencies (PIDs) are rare familial diseases that can be caused by a mutation in a variety of genes that affect the immune system, with currently 354 characterized monogenic forms (1). PIDs can affect the innate and adaptive arms of the immune system. Chronic granulomatous disease for example leaves phagocytes unable to produce microbicidal oxygen radicals, therefore severely compromising the innate immune system. PIDs that affect adaptive immunity usually exhibit functional defects of T and/or B lymphocytes as well as natural killer (NK) cells (2). Mutation in the genes *ZAP-70* and *LAT* for example lead to persistent infections that can be life-threatening due to defective T cell signaling. Some PIDs can be attributed to deficiencies in calcium (Ca^2+^) signaling (1). Ca^2+^ is a vital signaling molecule in all cells including immune cells and controls important processes like differentiation, homeostasis, activation, proliferation, and apoptosis (3). In lymphocytes, crosslinking the antigen receptor activates a signaling cascade that eventually leads to Ca^2+^ release from the endoplasmic reticulum (ER) into the cytoplasm (4). Upon Ca^2+^ depletion of the ER, Ca^2+^ channels in the plasma membrane open and a Ca^2+^ influx from the extracellular space is triggered. This process is called store-operated Ca^2+^ entry (SOCE) and the main plasma membrane channel involved in it is coined Ca^2+^ release-activated Ca^2+^ (CRAC) channel (5). The CRAC channel consists of the pore-forming unit called ORAI1 and a Ca^2+^ sensing protein named STIM1 that detects low levels of Ca^2+^ in the ER to activate the channel. Loss-of-function mutations in *ORAI1* or *STIM1* genes result in the partial abrogation of SOCE and defective T cell activation (6–9).

Apart from the CRAC channel however, there exist numerous other Ca^2+^ channels in the plasma membrane of lymphocytes that also contribute to the antigen receptor-mediated flux. Among them are the voltage-dependent Ca^2+^ channels (VDCCs), which have emerged as important players in immune cells (4). VDCCs consist of the pore-forming Cav (α1)-, the β regulatory-, and several other auxiliary subunits. They have been grouped into different families including the L-type Ca^2+^ channels, which are further divided into Cav1.1, 1.2, 1.3, and 1.4 (10). Since they are traditionally activated by a change in membrane potential, these channels have primarily been described in electrically excitable cells but more recent studies have also demonstrated that L-type Ca^2+^ channels play critical roles in murine and human leukocytes (10,11). Our lab has previously shown that Cav1.4 mRNA is subject to extensive splicing, including unique spliceforms in T lymphocytes resulting in the excision of the membrane-associated voltage sensor region resulting in the Cav1.4 channel becoming ligand gated (12). In another study with knock-out mice deficient in Cav1.4 our lab demonstrated that Cav1.4 plays an important role in T lymphocyte homeostasis and activation in mice. Specifically, Cav1.4 was required for the survival of naïve T lymphocytes as well as pathogen-specific T cell responses (13). Cav1.4, whose α1 subunit is encoded by the gene *CACNA1F* on the X chromosome, is also expressed in human T lymphocytes, which led us to hypothesize that it might be important for T cell function in humans as well (12,14).

For humans, the function of Cav1.4 is well described in the eye as its absence or reduced function can lead to incomplete congenital stationary night blindness (CSNB) (15). However, the role of Cav1.4 in the human immune system remains elusive. We had the opportunity to examine three brothers, each harboring the same *CACNA1F* mutation leading to incomplete CSNB, who we determined uniformly exhibit a PID. Here we describe the effect of Cav1.4 deficiency in human lymphocytes and mechanisms underlying a new X-linked immunological disorder.

## Results

### Case presentation

Three male siblings (16 – 20 years old) who each harbor the same mutation in the *CACNA1F* gene and who each present with immune deficiencies in addition to incomplete CSNB were studied. Their immune deficiencies include frequent recurrent infections, particularly of the ear and upper respiratory tract, tolerance defects, as seen by antinuclear antibodies in their serum (titer 1:160 for patient 1, 1:80 for patient 2, speckled). Other symptoms including aphthous ulcers, fatigue, muscle weakness, joint hypermobility, intermittent creatine phosphokinase (CPK) abnormalities, postural orthostatic tachycardia syndrome, Ehlers-Danlos syndrome and rashes were reported. A comprehensive list with all the patients’ conditions according to their mother can be found in ***Table S1***. A complete blood count with differential did not show any dramatic deviations from the normal ranges (***Table S2***). In previous clinical flow cytometric tests, it was found that one sibling exhibits a CD4/CD8 T cell ratio lower than the reference range while his brothers also had a ratio at the low end of the range. Additionally, two of the three siblings displayed a below normal frequency of naïve CD4+/CD45RA+ T cells (***Table S3***). Furthermore, two of the three siblings exhibited impaired responses to Tetanus, Pokeweed mitogen, and Candida, while responses to PHA and ConA were normal (***Table S4***). Finally, a high viral load of EBV and antibodies against EBV were detected in two of the three siblings (***Table S5***, third sibling not tested).

### Patients harbour R625X mutation in CACNA1F gene

The siblings as well as their mother underwent several genetic tests including Whole Exome Sequence Analysis by GeneDX (XomeDxPlus). In this analysis a R625X (p.Arg625Ter) point mutation that leads to a premature stop codon in the *CACNA1F* gene was discovered. The amino acid substitution is caused by the 1873C>T (NM_005183.3) nucleotide mutation changing the codon CGA to TGA in exon 14 in the voltage sensing domain of the protein (***Fig. 1A***). The dbSNP identifier is rs886039559 and the surrounding DNA sequence is GCAGCCCTTGGGCATCTCAGTGCTC[C/T]GATGTGTGCGCCTCCTCAGGATCTT. The R625X variant was previously described in a family with incomplete CSNB (16) and is predicted to cause loss of normal protein function due to protein truncation or nonsense-mediated mRNA decay. It was not observed in the approximately 6503 individuals in the NHLBI Exome Sequencing Project nor in the 60706 unrelated individuals in the ExAC database.

**Fig. 1.**
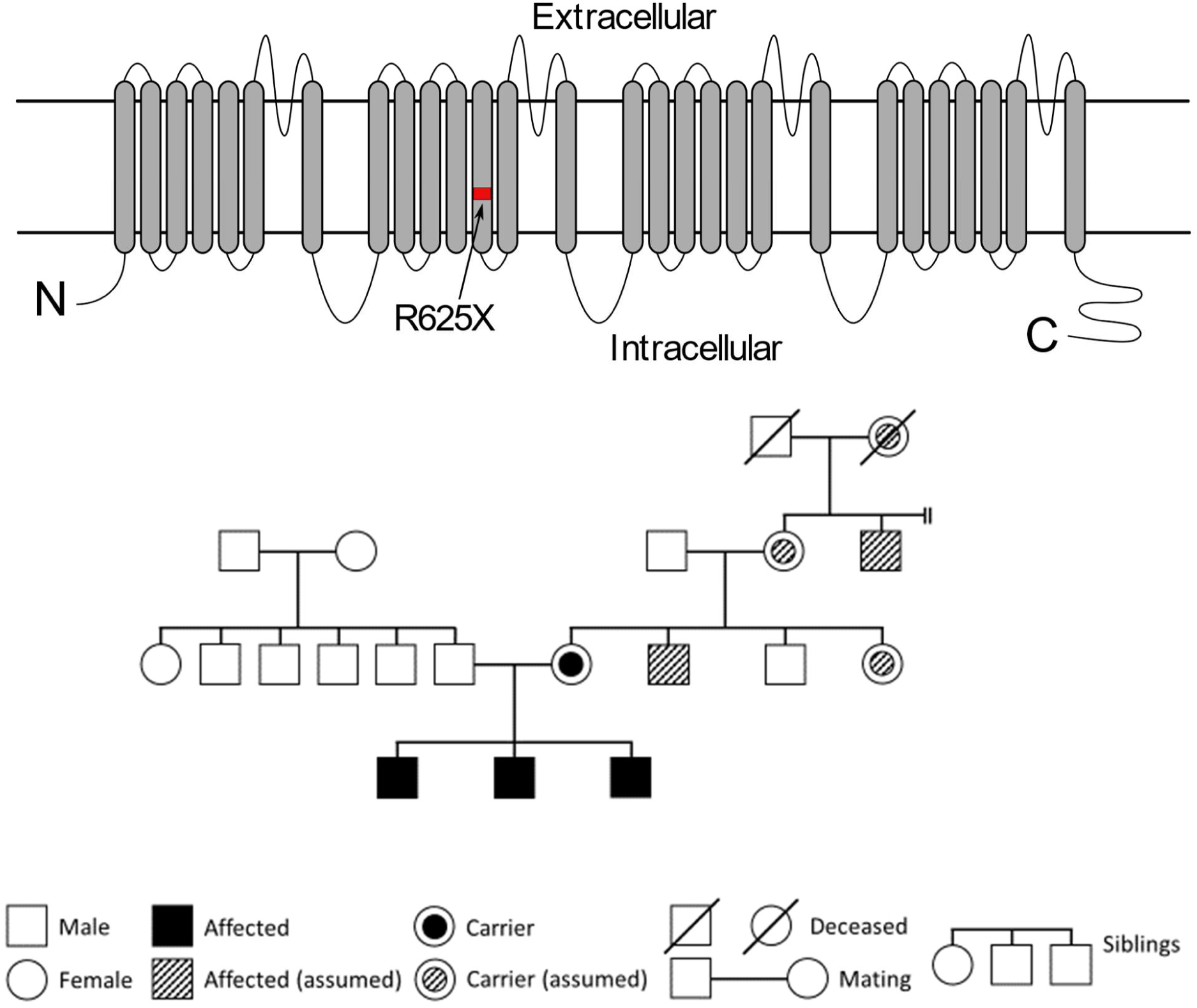
CACNA1F transmembrane protein cartoon and family pedigree of the three affected siblings. Cartoon depicting the structure of the α1 subunit of Cav1.4 and the location of the R625X mutation (A). The mother of the siblings is a confirmed carrier of the R625X mutation and other individuals on the maternal side of the family are assumed carriers of the allele. While the two affected (assumed) males display CSNB, the female carriers (assumed) exhibit autoimmune defects (B).

This variant is present in all three siblings who inherited the mutated allele from their mother who is a heterozygous carrier. Since *CACNA1F* is encoded on the X chromosome, the siblings are all hemizygous for the mutation. On the maternal side of the family there are two additional males who are also affected by CSNB and females including the siblings’ mother exhibit autoimmune defects. While their mother has been tested heterozygous for the R625X mutation the other individuals displaying CSNB and autoimmune defects are assumed carriers of the mutated allele (***Fig. 1B***).

The youngest of the three siblings is also hemizygous for the missense V40M variant in the *SH2D1A* gene (dbSNP rs199639961), which however did not change the expression of the encoded protein SAP on T and NK cells and is therefore not considered to be pathogenic. The middle sibling is homozygous and the oldest heterozygous for the Q45X (dbSNP rs17602729) and P81L (dbSNP rs61752479) variants in the *AMPD* gene. These *AMPD* variants have a high allele frequency and are only pathogenic in rare cases. The patients had other benign SNPs in genes *MEFV, NLRP3, TNFRSF1A* (rs224225, rs224224, rs224223, rs224213, rs224208, rs224207, rs224206, rs1231122, rs3806268, rs767455, rs1800693, rs12426675). An overview of the genetic variants identified by GeneDx can be found in ***Table S6***. The pathology in the three male patients is inherited as an X-linked genetic disease. The R625X mutation in *CACNA1F* is the only common pathogenic genetic alteration present in all three siblings. We therefore conclude that the R625X mutation in *CACNA1F* causes their immune dysfunction.

### Cav1.4 deficiency leads to a reduced CD4/CD8 T cell ratio

In order to further explore the role of Cav1.4 in humans, we isolated PBMCs from whole blood of these patients as well as several age- and sex-matched control donors. We analyzed the patients’ T cell populations by flow cytometry and could confirm the decreased CD4/CD8 T cell ratio found in previous clinical tests. This was due to a reduced frequency of CD4 T cells, which was compensated by an increase of CD4-CD8-double negative cells. The frequency of CD8 T cells and B cells (CD19+) was comparable to control donors (***Fig. 2A***).

**Fig. 2.**
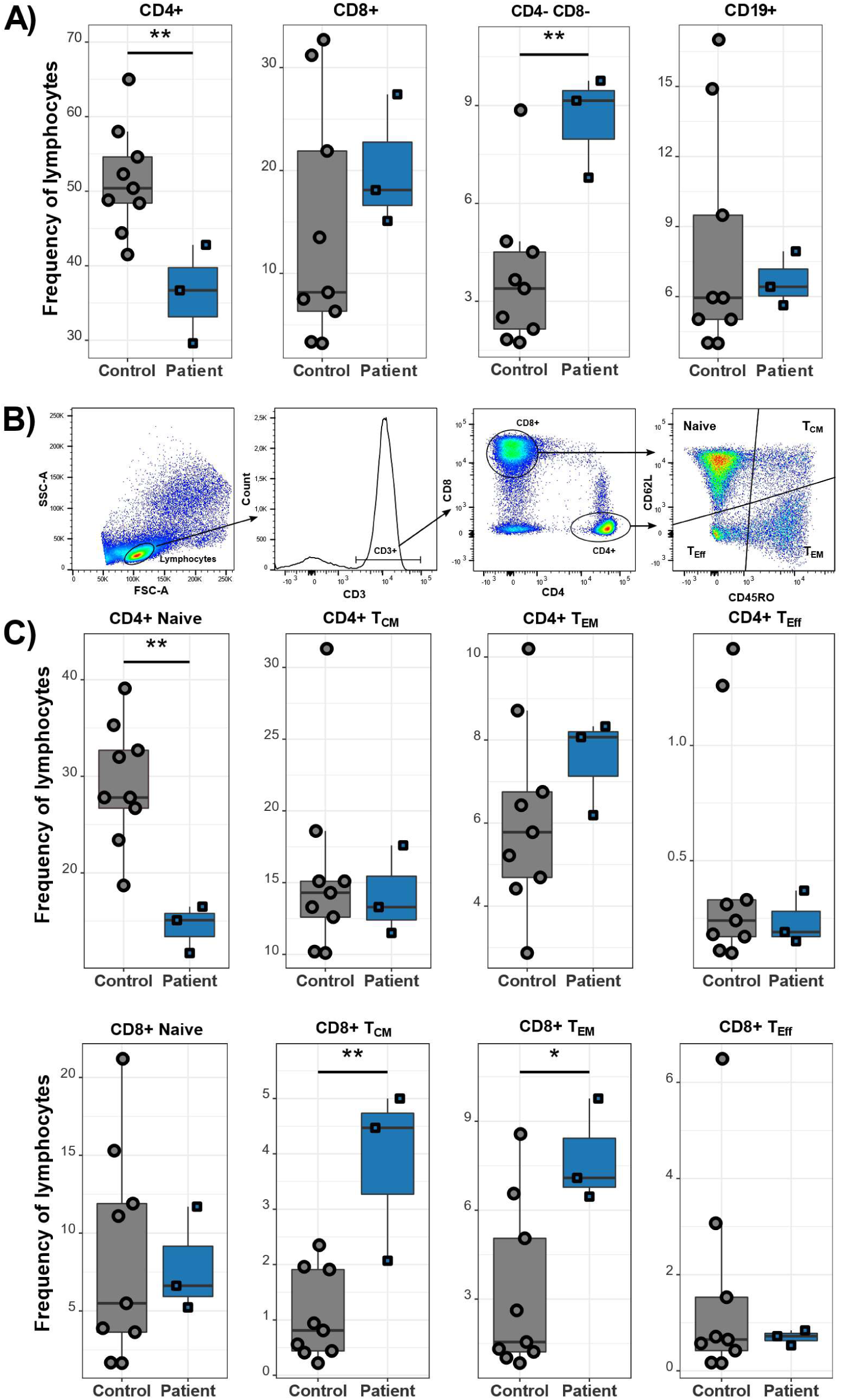
Cav1.4-deficient patients have a reduced number of CD4 T cells and an increased frequency of memory T cells. PBMCs of patients (n=3) and healthy controls (n=9) were stained with different antibodies and analyzed by flow cytometry. Frequencies of CD4+, CD8+, CD4-CD8- and CD19+ lymphocytes are shown (A). Gating strategy is shown (B). The different population frequencies shown in boxplots are classified as CD62L+ CD45RO- (naïve), CD62L+ CD45RO+ (T_CM_), CD62L-CD45RO+ (T_EM_), CD62L- CD45RO- (end-stage T_Eff_) (C). Representative of two technical replicates. *p<0.05 ** p<0.01.

### Cav1.4-deficient patients exhibit an increased frequency of memory and regulatory T cells

We next examined naïve and memory T cell subsets in these patients to assess to which specific population the reduced frequency of CD4 T cells can be attributed. In humans, naïve T cells are generally CD62L+, CCR7+, CD27+, CD45RA+, CD45RO-, central memory T cells (T_CM_) are CD62L+, CCR7+, CD27+, CD45RA-, CD45RO+, effector memory T cells (T_EM_) are CD62L-, CCR7-, CD27-, CD45RA-, CD45RO+ and end-stage effector T cells (T_Eff_) are CD62L-, CCR7-, CD27-, CD45RA+, CD45RO- (17,18). For this experiment, we used flow cytometry markers to group the cells into naïve (CD62L+ CD45RO-), T_CM_ (CD62L+ CD45RO+), T_EM_ (CD62L-CD45RO+), and end-stage T_Eff_ cells (CD62L-CD45RO-) (***Fig. 2B***). Compared to healthy control donors, the patients had a higher frequency of CD8 T_CM_ cells as well as an increased frequency of T_EM_ cells, which was more pronounced in the CD8 T cell subset. While their naïve CD8 T cell frequency was normal, the naïve CD4 T cell frequency was reduced (in agreement with previous clinical tests), which is the main reason for the reduced frequency of total CD4 T cells (***Fig. 2C***). Lastly, the patients also exhibited a significantly increased frequency of regulatory T lymphocytes as well as CD8 MAIT cells (***Fig. S1A/B***). The frequency of T lymphocytes expressing a γδ TCR was similar in patients and controls (***Fig. S1C***).

### T cells of Cav1.4-deficient patients have multiple upregulated inhibitory receptors

Next, we looked at the activation and exhaustion status of the patients’ T cells. T cell exhaustion develops during chronic infections as well as cancers and leads to poor effector function (19). Affected T cells can be identified by the continuous high expression levels of inhibitory receptors (IRs) like PD-1, CTLA-4, and TIGIT (*17*). Patients’ memory T and T_Eff_ cells of both the CD4 and CD8 subsets exhibited increased levels of PD-1 on their cell surface as seen by flow cytometry (***Fig. 3A***). T_CM_ cells and the CD4+ T_EM_ subset were especially affected. The patients’ PBMCs were also analyzed using CyTOF. Apart from confirming the expansion of T_CM_ and T_EM_ cells (not shown), we also observed the upregulation of multiple IRs that are consistent with CD8 T cell exhaustion. Apart from PD-1, other IRs including TIGIT and the transcription factors T-bet and EOMES, which are characteristic for exhausted T cells (20), were also significantly upregulated on the patients’ CD8 T cells. There was a significant increase in EOMES+ PD-1+ cells, which represent terminally differentiated exhausted T cells, that are known to have very limited proliferation and cytokine production potential (21) (***Fig. 3B***).

**Fig. 3.**
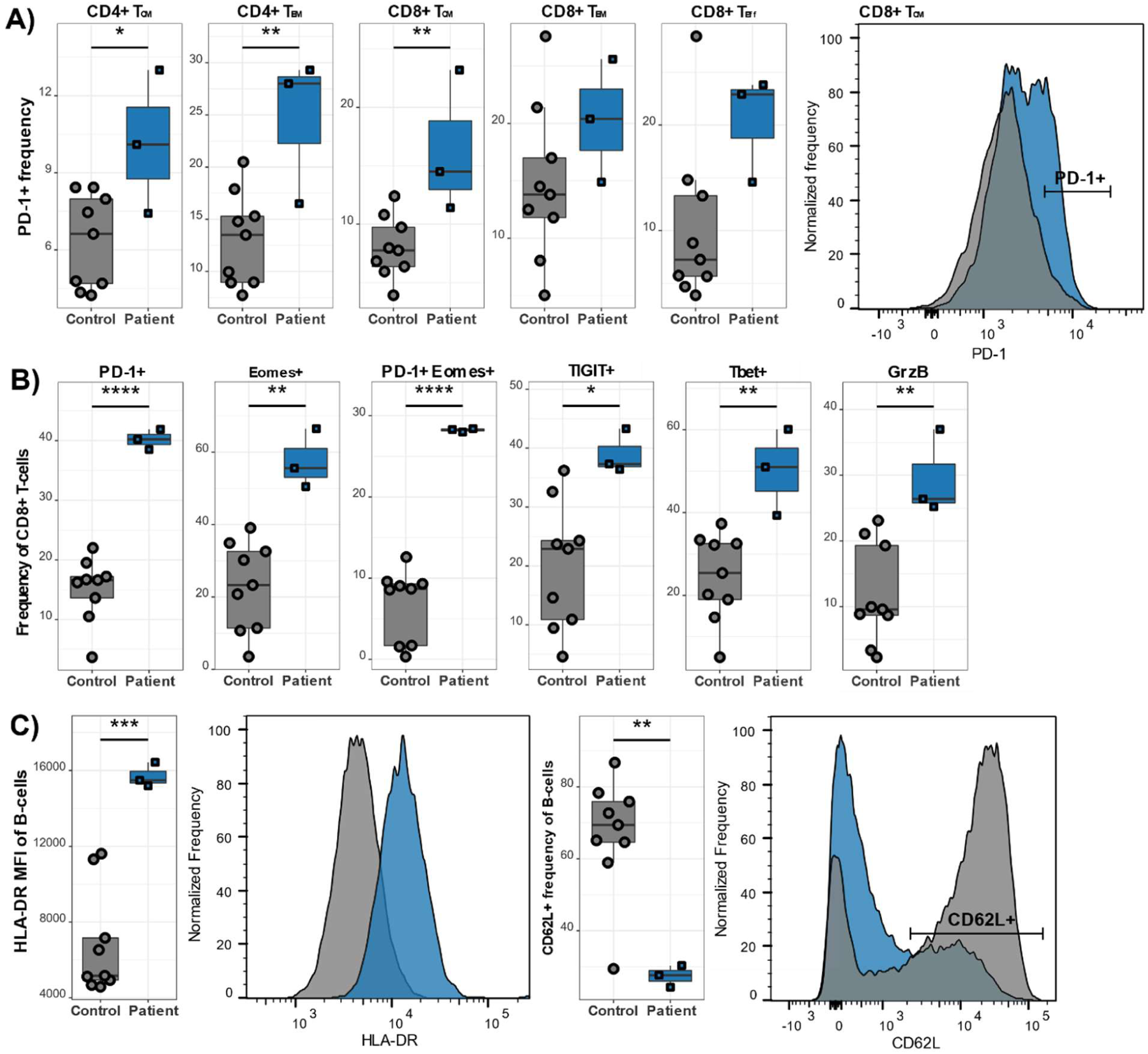
Cav1.4-deficient T cells show signs of exhaustion. B lymphocytes are chronically activated. PBMCs of patients (n=3) and healthy controls (n=9) were stained with different antibodies and analyzed by flow cytometry (A,C) / CyTOF (B). The PD-1+ frequency is shown in boxplots for the indicated populations, which are classified as CD62L+ CD45RO- (naïve), CD62L+ CD45RO+ (T_CM_), CD62L- CD45RO+ (T_EM_), CD62L- CD45RO- (end-stage T_Eff_) (A). Frequencies of CD8 T lymphocytes exhibiting other exhaustion markers are shown (B). The population quantified/shown in (C) is gated on B lymphocytes only (CD19+). Representative of two technical replicates (A,C). Experiment done once (B). Histograms show one representative sample of each genotype. * p<0.05 ** p<0.01 *** p<0.001 **** p<0.0001.

### B lymphocytes of Cav1.4-deficient patients are chronically activated

B lymphocytes are activated by binding to an antigen with their BCR. In T lymphocyte-dependent responses, they additionally bind to CD40L, expressed on T helper lymphocytes, which together with secreted cytokines like IL-4 and IL-21 provides a costimulatory signal for the B lymphocytes (22). Throughout this process, B lymphocytes upregulate several activation markers on their cell surface, including MHCII. B cells (CD19+) of the patients expressed high levels of the MHCII isotype HLA-DR (***Fig. 3C***), implying that they are in an activated state. Also, the expression levels of the adhesion molecule CD62L, which is shed upon B lymphocyte activation, were strongly reduced on the surface of the patients’ B lymphocytes. Additionally, the patients exhibited a higher frequency of class-switched B cells (***Fig. S1D***).

### Cav1.4 deficiency impairs Ca^2+^ flux in T lymphocytes but not B lymphocytes

We next performed a Ca^2+^ flux assay using the patients’ PBMCs. PBMCs were labeled with Ca^2+^ dyes and different lymphocyte markers, stimulated with thapsigargin, and the Ca^2+^ flux was recorded by flow cytometry. While B lymphocytes of the patients displayed a normal flux, the Ca^2+^ mobilization in their CD8 and CD4 T lymphocytes was reduced, compared to healthy donor cells (***Fig. 4***). Thapsigargin blocks the reuptake of Ca^2+^ into the ER, which triggers SOCE without engaging the antigen receptor and the associated proximal TCR/BCR signaling pathway. We therefore hypothesize that Cav1.4 is also involved during SOCE and that its absence leads to a reduction of the induced Ca^2+^ flux. Interestingly, stimulation with α-CD3 triggered a similar Ca^2+^ flux in CD4 and CD8 T cells of Cav1.4-deficient patients and healthy controls (not shown). We also tested whether the shRNA-mediated knockdown of Cav1.4 in Jurkat T cells reduced the TCR-induced Ca^2+^ flux. The shRNA treated cells, which exhibited an 80% reduction of *Cacna1f* mRNA, demonstrated a reduced Ca^2+^ flux upon α-CD3 stimulation (***Fig. S2***).

**Fig. 4.**
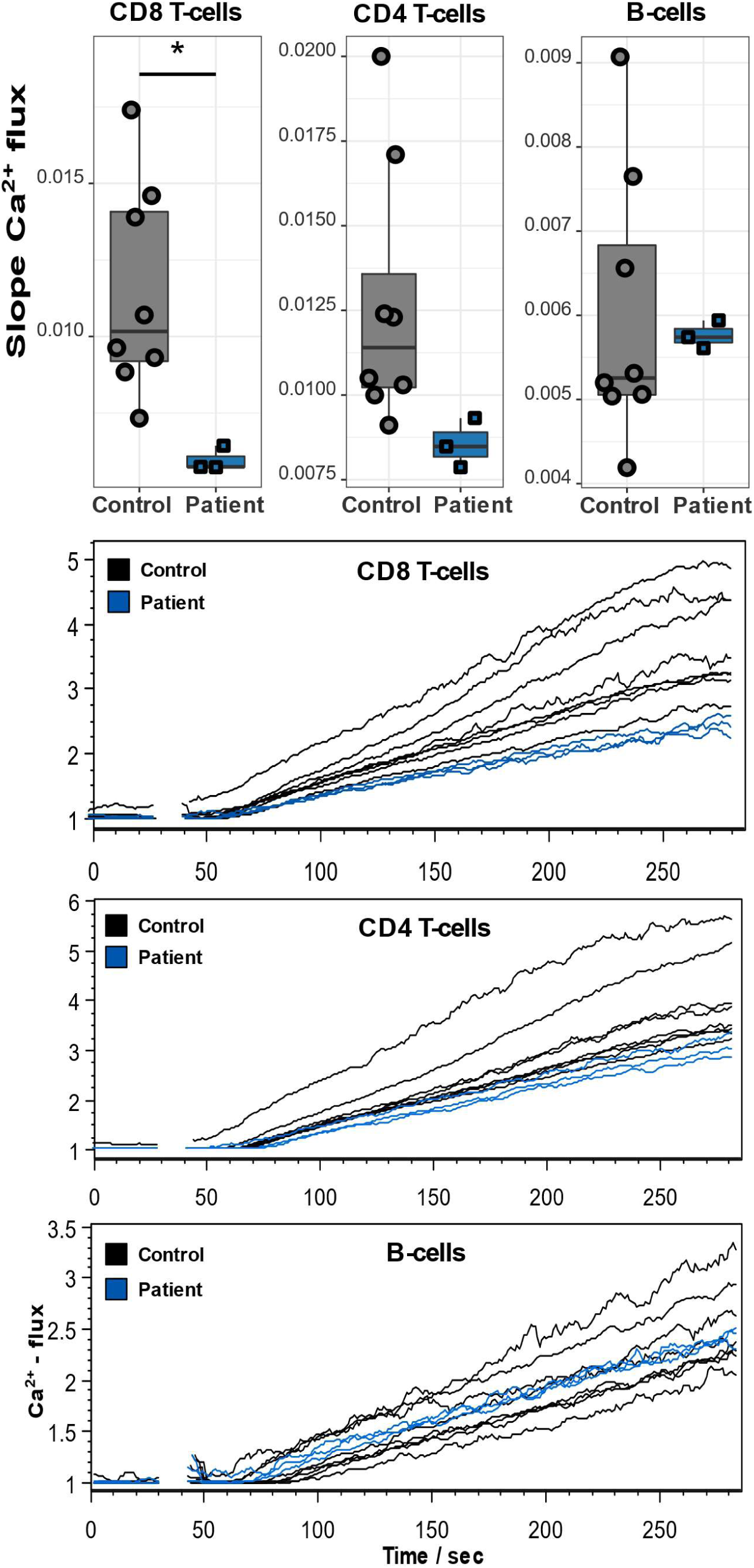
Cav1.4-deficient T lymphocytes exhibit a reduced Ca^2+^ flux. PBMCs of patients (n=3) and healthy controls (n=8) were stained with Ca^2+^ dyes and different T and B lymphocyte-specific antibodies and analyzed by flow cytometry. Thapsigargin was added to cells after 30 seconds of acquisition. The boxplots show the quantified slopes of increasing Ca^2+^ concentration for each cell type (A). The flow cytometry kinetics plots show the actual Ca^2+^ influx over time (B). Representative of two technical replicates. * p<0.05.

### Cytokine secretion in Cav1.4-deficient T lymphocytes is comparable to healthy donor levels

Lastly, to evaluate T lymphocyte function, we stimulated PBMCs with α-CD3 and α-CD28/49d and then quantified the expression of the T lymphocyte activation marker CD38 as well as the intracellular cytokine levels of IFN-γ and IL-2. Contrary to our expectations they were upregulated similarly in patients’ and controls’ non-naive T lymphocytes (gated by excluding CD27+ CD45RA+) (***Fig. S3***). In fact, the patients’ T lymphocytes had an increased frequency of CD38+ T lymphocytes compared to the controls after stimulation. However, it is important to note that the presence of exhaustion does not preclude the potential to secrete cytokines. In comparison to mouse studies, which are generally based on comparison of the activation state and function of antigen specific lymphocytes in a specific infection, in human studies we are often unable to focus on an antigen specific sub-population for appropriate comparison of functional capacity between affected and control subjects.

## Discussion

We have characterized the first example of a mutated Cav channel causing a primary immunodeficiency in humans, which is inherited as an X-linked genetic disease. The patients we examined have an R625X point mutation that leads to a premature stop codon in their *CACNA1F* gene. No other pathogenic SNPs exist in any of the patients. Thus, the R625X mutation in *CACNA1F* is the only common pathogenic genetic alteration present in all three siblings. We therefore concluded that the R625X mutation in *CACNA1F* causes their immune dysfunction. Many mutations in *CACNA1F*, including this one, have been linked to CSNB (16). The patients in the present study also exhibit this phenotype and participated in a study further describing their disease (23). This, however, is the first report of an immune disorder caused by a mutation in the *CACNA1F* gene, and appears to be a novel phenotype amongst all human primary immunodeficiencies.

The siblings exhibit a low CD4 T lymphocyte frequency and hence a low CD4/CD8 T lymphocyte ratio and also present a high frequency of dysfunctional T lymphocytes and an expansion of memory T lymphocytes. Additionally, the frequencies of regulator T lymphocytes as well as CD8 MAIT lymphocytes are significantly elevated in the siblings. The B lymphocytes of Cav1.4-deficient patients were chronically activated, as indicated by the upregulation of several activation markers. Interestingly, the human *CACNA1F*-/-B lymphocytes also exhibited a loss of CD62L. CD62L is an adhesion molecule that is necessary for cell migration to the site of inflammation and does so by mediating the rolling of leukocytes on the endothelium. It was found that the P2X7 receptor agonist benzoyl-benzoyl-ATP induces shedding of CD62L on lymphocytes (24). As P2X receptors also flux Ca^2+^ across the plasma membrane of lymphocytes, this implies a potential role of Ca^2+^ in the shedding of CD62L. I nterestingly, phorbol 12-myristate 13-acetate, a strong B cell activator, also induced shedding of CD62L in B-CLL lymphocytes (25), which is why we also considered CD62L downregulation as a marker of B lymphocyte activation. Due to the unchanged Ca^2+^ flux upon thapsigargin treatment however, we hypothesize that the chronic B lymphocyte activation is an indirect effect of the R625X mutation in *CACNA1F*.

CRAC channel deficiencies do not impair T lymphocyte development but severely reduce proliferation and effector function due to strongly impaired SOCE (26–28). Thapsigargin treatment induced a lower Ca^2+^ influx into our patients’ T lymphocytes than into those of healthy controls. Since thapsigargin skips other TCR/BCR-induced signaling pathways and directly induces SOCE, this reduction in Ca^2+^ mobilization demonstrates that, like the CRAC channel, Cav1.4 also contributes to SOCE. However, when stimulating T lymphocytes via the TCR using α-CD3, the differences in Ca^2+^ flux were not observed. On the other hand, Jurkat T cells treated with Cav1.4 shRNA did show a marked reduction of α-CD3-induced Ca^2+^ flux. Also, we have previously demonstrated that the L-type Ca^2+^ channel inhibitor nifedipine blocks a TCR-induced Ca^2+^ flux in Jurkat T cells as well as human PBMCs (14). We therefore hypothesize that the reason we did not observe a reduced TCR-induced Ca^2+^ flux in our patients may be that their T lymphocyte subset composition is different from the healthy controls we compare them to, as shown in ***Fig. 1C***. Indeed, it has previously been demonstrated that CD45RO+ memory T cells exhibit a higher α-CD3-mediated Ca^2+^ response than CD45RA+ naïve T lymphocytes (29). The higher frequency of central and effector memory T lymphocytes in our patients might therefore make up for the expected reduced Ca^2+^ flux when averaging the Ca^2+^ signal of naïve and memory T lymphocytes together. Thapsigargin simulation on the other hand, shows a reduced Ca^2+^ flux even when comparing the mixed T lymphocyte populations, since it bypasses the proximal TCR signaling pathway where the different signaling strengths of naïve and memory T lymphocytes most likely manifest.

We have previously demonstrated that despite diminishing the Ca^2+^ flux nifedipine treatment also reduced the TCR-induced translocation of NFAT, activity of ERK1/2 as well as the secretion of IL-2 and the expression of the IL-2 receptor (14). The retained ability to secrete cytokines in our patient’s T lymphocytes is not necessarily inappropriate for exhausted T lymphocytes; exhaustion does not yield lack of function and in human patients when studying non-antigen specific responses, it can be challenging to detect the altered function comparing to healthy controls. Studies have shown that the overexpression of IRs does not always correlate with decreased cytokine secretion capacity as often seen in exhaustion (30). Instead, the differentiation and activation status of the T lymphocytes is concomitant with the expression of IRs (30–33). However, previous clinical tests demonstrating diminished lymphocyte proliferation in response to antigenic as well as mitogenic stimuli (**Table S4**) do indeed suggest functional exhaustion of the patients’ T cells.

Also in mice, Cav1.4-deficient T lymphocytes exhibited reduced thapsigargin-induced Ca^2+^ mobilization, as shown in previous work in our lab (13). In β3 KO CD4 T lymphocytes, CD3 crosslinking also led to a diminished Ca^2+^ flux but, conversely, thapsigargin-induced Ca^2+^ flux was normal (34). Despite these discrepancies, both KO mouse models displayed impaired nuclear translocation of NFAT, which resulted in reduced cytokine production (13,34,35).

The three siblings incur frequent recurrent ear-as well as upper respiratory tract infections. Previous clinical tests have shown that T lymphocyte proliferation of the patients is decreased in response to antigens and mitogens, which indeed suggests that Cav1.4-deficient lymphocytes (despite normal TCR-induced cytokine responses) cannot efficiently respond to pathogens and clear infections. In the patients, this would also provide an explanation for their phenotype: the recurrent infections lead to the increase in memory T lymphocyte frequency and upregulation of exhaustion markers as well as the expansion of regulatory T lymphocytes. Also, MAIT lymphocytes preoccupied with recognizing vitamin-B metabolites from bacteria and their expansion implies the patients had all experienced chronically bacterial infections. Because of these frequent infections, the B lymphocytes of the patients also remain in a chronic state of activation. Alternatively, the phenotype might also arise independent of an underlying infection and instead result from a lymphocyte development defect due to Cav1.4 deficiency. This is supported by the fact that the patients have not reported any infections immediately prior to our analysis and the same phenotype was found using samples drawn at two different time points (data only shown for one). Lastly, the increase of anti-nuclear antibodies that was found in the three brothers suggests that Cav1.4 deficiency also might have an impact on tolerance. Abnormal T lymphocytes selection would also provide another explanation for the upregulation of the different activation/exhaustion markers on T lymphocytes.

It is important to note that besides the many cases of CSNB that are caused by mutations in *CACNA1F*, a growing list of patients do complain about recurrent infections and therefore and seem to display the immune phenotype we observed. Several affected families reporting immune-related symptoms have self-founded a Facebook group (https://www.facebook.com/groups/883482391791546/) in which they share their experiences of living with this gene defect. It is possible that a specific, potentially chronic infection triggered the disease in our patients. In many PIDs EBV as well as cytomegalovirus infections are reported to exacerbate the disease. An example for this is XLP disorder where an EBV infection leads to disease progression resulting in hyperproliferation of T lymphocytes and an aggravated cellular immune response. Interestingly, in previous clinical tests two of our patients (third patient not tested) were shown to have high levels of serum antibodies against EBV as well as high EBV viral loads (**Table S5**) and they reported that their symptoms worsened after mononucleosis during their teens. This deteriorating of a Cav1.4-related condition is also supported by a recent publication of our lab, which concludes that murine gamma herpesvirus 68, a viral ortholog of human EBV, can exacerbate the phenotype of a Cav1.4-deficient mouse model (36).

The severity of a phenotype in patients can also be affected by modifier genes. SNPs in modifier genes can alter phenotypes that are usually caused by a mutation in another gene, the so called target gene (37). This can affect the penetrance and expressivity of the target gene mutation. Modifier genes have been proposed to alter disease severity of night blindness of patient with a *CACNA1F* mutation (38). It is therefore plausible that modifier alleles also alter the immune phenotype caused by Cav1.4 deficiency and thereby change the penetrance and expressivity of the disease. An incomplete penetrance can also be observed in STIM1 deficiency where two cousins that, despite reduced SOCE and impaired NK effector functions, lack clinical symptoms (39). Lastly, following our original description of alternative splicing in Cav genes and the existence of spliceforms of Cav1.4 mRNA unique to T lymphocytes (12), at least nineteen, naturally occurring alternative splice variants of the native Cav1.4 mRNA were detected in the retina (40). It is possible that specific mutations in *CACNA1F* may be spliced out resulting in partial phenotypes depending on the spliceforms that remain unaffected.

In conclusion, we have identified the first primary human immunodeficiency caused by a genetic mutation in any L-type Ca^2+^ channel. This mutation results in impaired Ca^2+^ signaling in T cells and causes T lymphocyte dysfunction as seen in chronic infections and cancer with exhaustion in the setting of chronic antigen exposure. These studies establish the importance of L-type Ca^2+^ channels to immune physiology in humans.

## Methods

### Patient recruitment

CHOP: Patients were enrolled on a CHOP IRB approved protocol (PIs: Kathleen Sullivan or Neil Romberg). Penn: Patients were enrolled on a UPenn approved protocol (PI: Ramin Herati) or via the Human Immunology Core. UBC: Clinical Research Ethics Board granted approval for the human study (PI:Wilf Jefferies). Informed consent was obtained from all volunteers before whole-blood donation.

### PBMC acquisition

for CyTOF and cytokine analysis: Venous blood was obtained by venipuncture and prepared for peripheral blood mononuclear cells (PBMCs) using the SepMate system (STEMCELL Technologies) which were cryopreserved until used for the following studies.

For lymphocyte population flow cytometry and Ca^2+^ flux: 10 ml of whole blood were collected into tubes containing sodium citrate (12 mM). PBMCs were then isolated using Lymphoprep (STEMCELL Technologies, cat. nr. 07801) according to protocol and used in subsequent experiment (lymphocyte population analysis) or frozen down in fetal bovine serum (FBS) with 10% DMSO (Thermo Fisher) for later use (Ca^2+^ flux).

### Flow cytometry human lymphocyte population analysis

Freshly isolated or cryopreserved 2 × 10^6^ human PBMCs, resuspended in PBS with 2% FBS, were labelled for 30 minutes at 4°C in the dark with different antibody panels. The antibodies used were: α-CD8 (RPA-T8, BioLegend, cat. nr. 301005), α-CD4 (RPA-T4, BioLegend, cat. nr. 300511), α-CD3 (UCHT1, BioLegend, cat. nr. 300427), α-CD62L (DREG-56, BioLegend, cat. nr. 304841), α-CD45RO (UCHL1, BioLegend, cat. nr. 304205), α-PD-1 (EH12.2H7, BioLegend, cat. nr. 329915), α-IL-7R (A019D5, BioLegend, cat. nr. 351317), α-CD19 (HIB19, BioLegend, cat. nr. 302211), α-PD-L1 (29E.2A3, BioLegend, cat. nr. 329717), α-HLA-DR (L243, BioLegend, cat. nr. 307623), α-CD32 (FUN-2, BioLegend, cat. nr. 303205), α-CD14 (63D3, BioLegend, cat. nr. 367111). The cells were then washed twice with PBS, resuspended in PBS with 2% FBS and data were acquired on an LSRII (BD Biosciences) and analysed with FlowJo v10 software (Treestar, Inc).

### Mass cytometry (CyTOF)

Reagents for mass cytometry were purchased or generated using MAXPAR kit (Fluidigm) to custom conjugate using isotope-loaded polymers. Mass cytometry antibodies in this study are shown in ***Table S7***. Staining was performed as previously published (41). Single cell suspensions of PBMCs were thawed and incubated with 20μM Lanthanum-139 (Trace Sciences)-loaded maleimido-mono-amide-DOTA (Macrocyclics) in PBS 10 minutes to allow live/dead cell discrimination. Cells were washed in staining buffer and then resuspended in master mix of surface antibodies, incubated for 30 minutes at room temperature and washed twice in staining buffer. Cells were then fixed and permeabilized using the FoxP3 staining buffer kit (eBioscience) and then washed twice with 1X Permeabilization Buffer (eBioscience). Cells were then stained intracellularly for 60 minutes at room temperature. Three more washes were performed in 1X Permeabilization Buffer before being fixed using 1.6% PFA (Electron Microscopy Sciences) solution with Iridium and Osmium overnight at 4 degrees Celsius. Cells were washed twice in PBS and once in dH20 prior to data acquisition on a CyTOF Helios (Fluidigm). We used bead-based normalization of CyTOF data using the Nolan lab scripts (https://github/com/nolanlab/bead-normalization/releases). FSC files were analyzed using FlowJo v10 (Treestar, Inc) and viSNE (Cytobank).

### Flow cytometry for human cytokine analysis

Cells were stimulated with α-CD3 (coated plates, OKT3) and α-CD28/CD49d (soluble, L293, L25, BD Biosciences) for 4.5 hours, in the presence of brefeldin A (BD Biosciences) and monensin (BD Biosciences). Cells were then stained with cell surface and intracellular antibodies for flow cytometry. Flow cytometry antibodies in this study are shown in ***Table S8***.

### Ca^2+^ flux with human PBMCs

Frozen 4 × 10^6^ human PBMCs were thawed, resuspended in HBSS with 2% FBS, and labelled with 1 μM Fluo-4, 2 μM Fura Red (Thermo Fisher, cat. nr. F14201, F3021) for 45 minutes at room temperature in the dark. Cells were then washed, stained with α-CD19 (HIB19, BioLegend, cat. nr. 302211), α-CD8 (RPA-T8, BioLegend, cat. nr. 301026), α-CD4 (RPA-T4, BioLegend, cat. nr. 300511) for 30 minutes on ice, washed again, and resuspended in HBSS with 2% FBS. After prewarming the cells for 15 minutes at 37°C, baseline Ca2+ levels were acquired for 30 seconds by flow cytometry. 1 μM thapsigargin (Thermo Fisher, cat. nr. T7458) was then added to the cells and acquisition was continued for a total of 5 minutes. Data were acquired on an LSRII (BD Biosciences) and analysed with FlowJo v9 software (Treestar, Inc).

### shRNA transfection of Jurkat T cells

pGFP-V-RS vector (OriGene Technologies) expressing shRNA targeting human *CACNA1F* gene or pGFP-V-RS vector with scrambled shRNA cassette (control) was transfected into Jurkat T cells by Neon electroporation system (Invitrogen) according to manufacturer’s guidelines. shRNA targeting sequence for CACNA1F is CCTGCACATAGTGCTCAATTCCATCATGA. Knockdown efficiency of the CACNA1F gene expression was determined by RT-PCR.

### Ca^2+^ flux with Jurkat T cells

10^6^ transfected (scrambled or Cav1.4 shRNA) Jurkat T cells were resuspended in HBSS with 2% FBS, and labelled with 1 μM Fluo-4, 2 μM Fura Red (Thermo Fisher, cat. nr. F14201, F3021) for 45 minutes at room temperature in the dark. Cells were then washed and again resuspended in HBSS with 2% FBS. After prewarming the cells for 15 minutes at 37°C, baseline Ca^2+^ levels were acquired for 30 seconds by flow cytometry. α-CD3 (1:1000 dilution, HIT3a, Invitrogen) was then added to the cells and acquisition was continued for a total of 10 minutes. Data were acquired on an LSRII (BD Biosciences) and analysed with FlowJo v9 software (Treestar, Inc).

### Exome Sequencing by GeneDX

Using genomic DNA from specimen of the three siblings and their mother, the Agilent Clinical Research Exome kit was used to target the exonic regions and flanking splice junctions of the genome. These targeted regions were sequenced simultaneously by massively parallel (NextGen) sequencing on an Illumina HiSeq sequencing system with 100bp paired-end reads. Bi-directional sequence was assembled, aligned to reference gene sequences based on human genome build GRCh37/UCSC hg19, and analysed for sequence variants using a custom-developed analysis tool (Xome Analyzer, GeneDx, Gaithersburg, MD, USA). Capillary sequencing or another appropriate method was used to confirm all potentially pathogenic variants identified in the siblings and their parents. Sequence alterations were reported according to the Human Genome Variation Society (HGVS) nomenclature guidelines. The exome was covered at a mean depth of 138x, with a quality threshold of 93.7%.

### Sample size estimation

Sample size of the patient group was limited to three as we were unable to recruit more patients with a *CACNA1F* mutation for this study. The sample size of the control group was estimated to detect a Cohen’s effect size of d = 2 with a significance level of α = 0.05 and power of 75%. This results into a sample size of n ≈ 8.1 for the control group and was calculated using R’s pwr.t2n.test() function.

### Statistical tests

Statistical significance was determined with 2-sided unpaired Student’s t tests using R’s t.test() function. Technical replicates are from the same blood sample but different aliquots (either freshly isolated or cryopreserved PBMCs), analyzed on different days.

### Boxplots

Points represent the individual data measurements. The center line in the boxplot is the median while the box represents the interquartile range (IQR). The upper and lower limits of the box represent the first (Q1) and the third quartile (Q3). The upper whisker limit is the max value < (Q3 + 1.5 * IQR). The lower whisker limit is the min value > (Q1 - 1.5 * IQR). Points that lie outside the whiskers range are outliers.

### Data Sharing Statement

For original data, please contact.

## Authorship Contributions

### Conceived of the Study

WAJ

### Designed research

FF, SRS, CGP, SEH, KES, WAJ

### Performed research

FF, SRS, CJL, SEH, OK, KCOB, KM, MR, RSH, NDR, EJW, KES,

### Analyzed data

all authors

### Wrote paper

FF, WAJ

### Edited paper

all authors

## Acknowledgments

We would like to thank the patients and their family for their willingness to participate in this study. We thank Woosuk S. Hur and Dr. Christian Kastrup for providing healthy control blood samples. We also thank our sponsors for their funding for this work: F.F. was the recipient of a DOC Fellowship of the Austrian Academy of Science and the Dmitry Apel Memorial Scholarship; S.R.S. was the recipient of a John Richard Turner Fellowship; H.A. was the recipient of a Centre for Blood Research Graduate Student Award; W.A.J. was supported by grants from the Canadian Institutes of Health Research (IPR-139079; MOP-102698), and a grant from Pascal Biosciences, Inc.;. E.J.W. was supported by the Parker Institute for Cancer Immunotherapy.

## Supplementary Materials

**Fig. S1.**
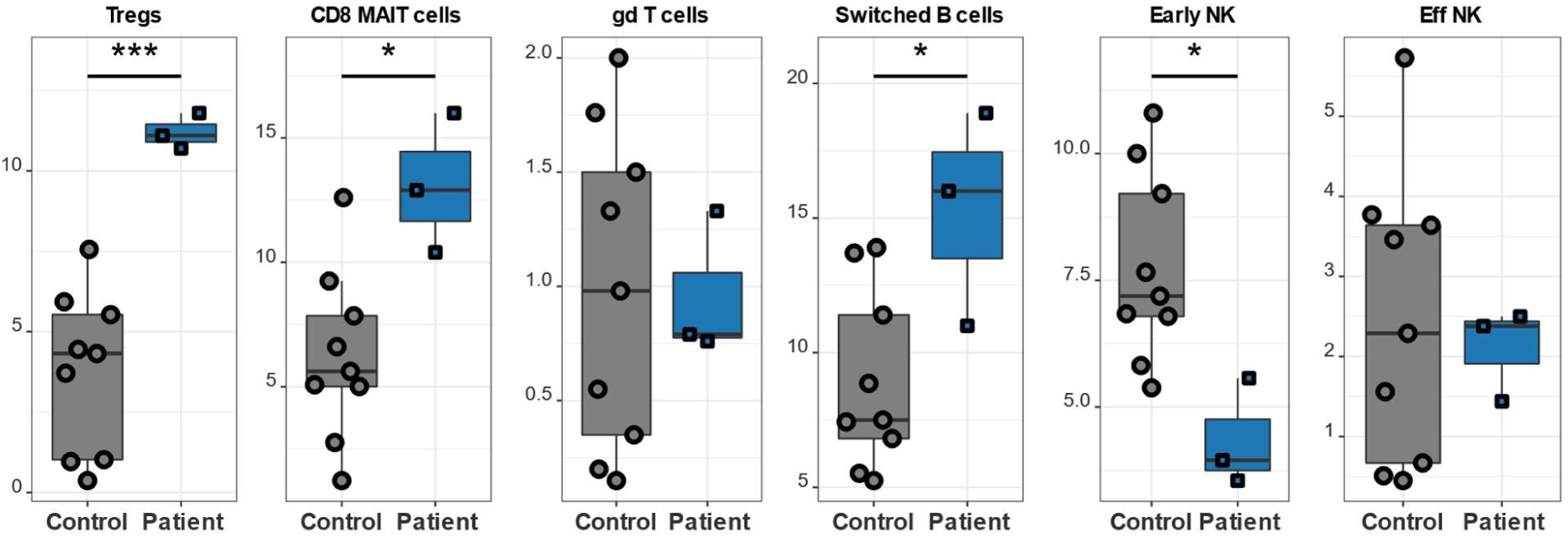
PBMCs of patients (n=3) and healthy controls (n=9) were stained with different antibodies and analyzed by CyTOF. This experiment was only done once.

**Fig. S2.**
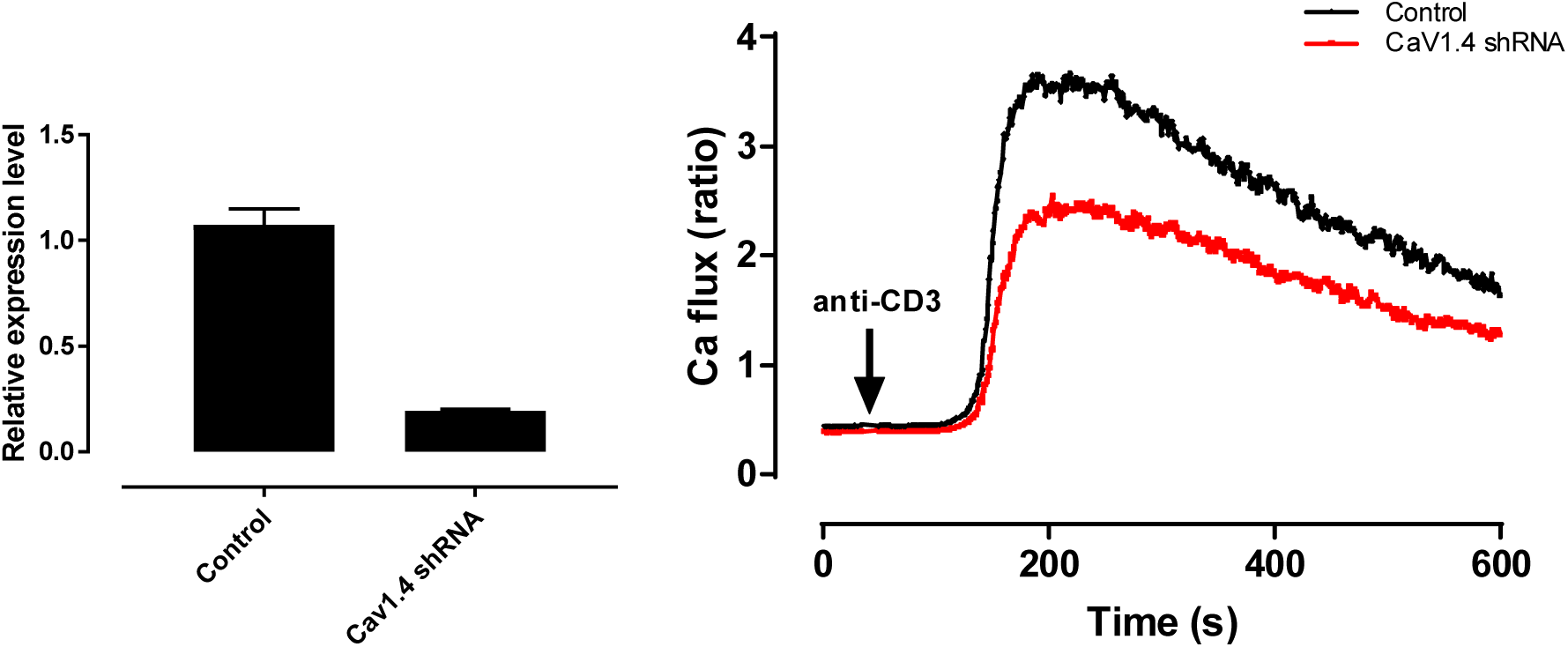
Cav1.4 shRNA treated Jurkat T cells exhibit a reduced TCR-induced Ca^2+^ flux. (A) Cav1.4 gene expression by qRT-PCR in Jurkat cells after 48 hours of Cav1.4 shRNA transfection. (B) Jurkat T cells, transfected with either Cav1.4 shRNA or scrambled control shRNA were loaded with Ca^2+^ dyes and analyzed by flow cytometry. After 30 seconds the cells were activated with α-CD3. Knockdown was performed once, Ca^2+^ flux experiment is a representative of two technical replicates.

**Fig. S3.**
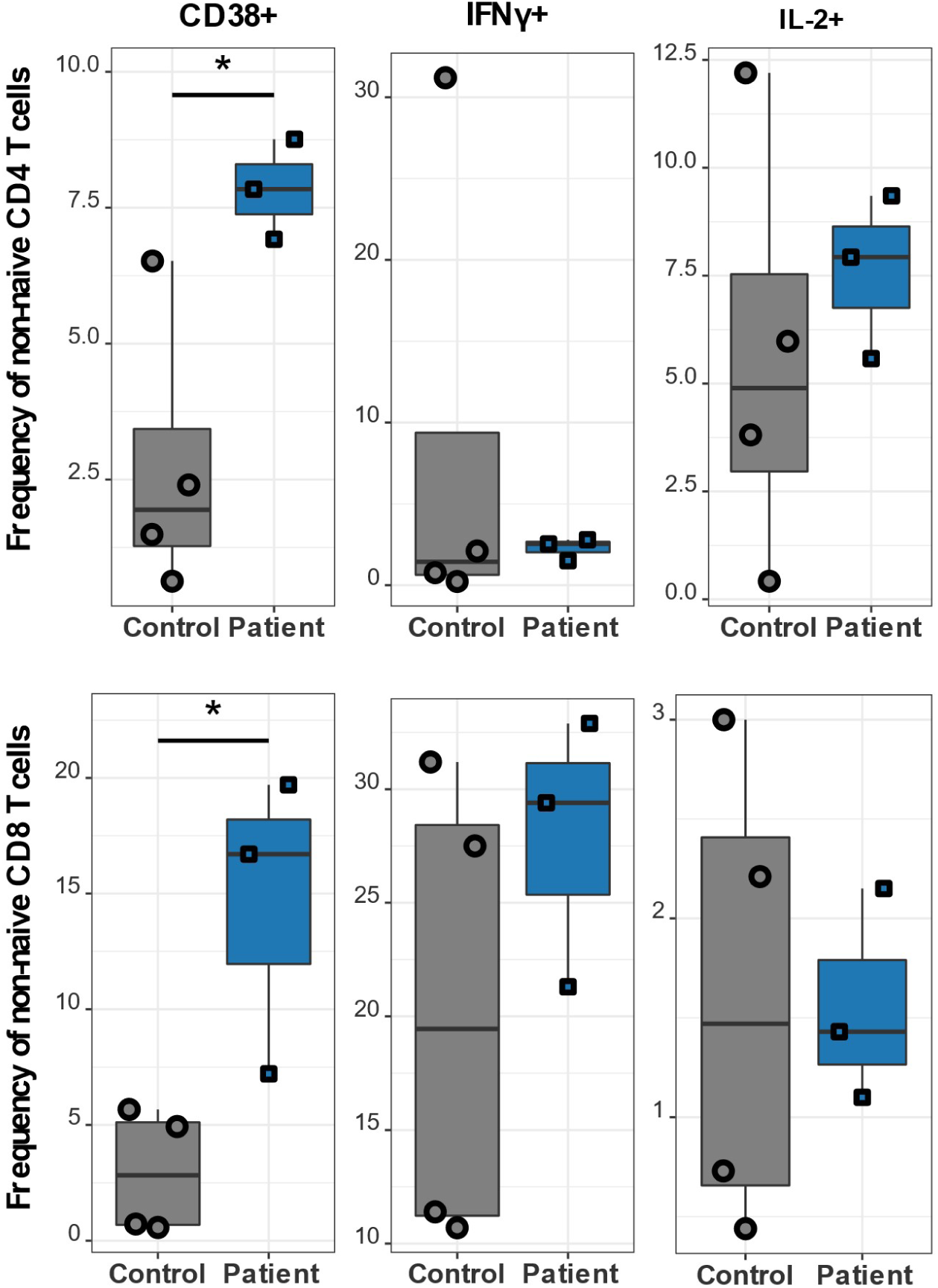
T lymphocyte activation marker CD38 is increased, while cytokine secretion is unchanged in Cav1.4-deficient patients upon stimulation. PBMCs of patients (n=3) and healthy donors (n=4) were stimulated with α-CD3 and α-CD28/49d for 4.5 hours and subsequently stained with various antibodies for extracellular markers and intracellular cytokines. The frequencies of CD38+, IFNγ+, and IL-2+ lymphocytes are shown for CD4 and CD8 non-naïve T lymphocytes, which are classified as NOT CD27+ CD45RA+. This experiment was only done once.

**Table S1.** Clinical history of patients.

|  | Patient 1 | Patient 2 | Patient 3 |
| --- | --- | --- | --- |
| **Lab tests** |  |  |  |
| Impaired lymphocyte proliferation in response to antigens and mitogens | X | X |  |
| Polycythemia | X | X | X |
| **Skin conditions** |  |  |  |
| Rash - dermatomyositis pattern |  |  | X |
| Rashes - Urticaria | X |  |  |
| Dermatographism | X |  |  |
| **Eye disorders** |  |  |  |
| Optic atrophy | X | X | X |
| Congenital stationary night blindness | X | X | X |
| Dry Eye Syndrome |  |  | X |
| Keratitis |  |  | X |
| Blepharitis |  |  | X |
| Conjunctivitis |  |  | X |
| **Muscle / joint disorders** |  |  |  |
| Myalgia | X |  | X |
| Arthralgia | X |  | X |
| Muscle cramps |  |  | X |
| Bilateral knee joint effusions |  |  | X |
| Muscle weakness | X |  |  |
| **Ear, nose, and throat disorders** |  |  |  |
| Chronic ear infections |  |  | X |
| Chronic rhinitis | X |  |  |
| Reoccurring pharyngitis | X |  |  |
| Sinusitis |  | X |  |
| **Infections** |  |  |  |
| Pneumonia | X | X | X |
| Frequent viral infections | X | X | X |
| Recurring EBV |  | X | X |
| **Other features** |  |  |  |
| Bilateral hydroceles |  |  | X |
| Torsion of the appendix testis |  |  | X |
| Testicular pain | X |  |  |
| Gastroesophageal junction polyps |  |  | X |
| Gastritis | X | X | X |
| Intussusception |  | X |  |
| Aphthous ulcers | X | X | X |
| Fatigue and malaise | X | X | X |
| Undifferentiated connective tissue disease |  |  | X |
| Kidney stones |  |  | X |
| Raynauds phenomenon | X |  |  |
| Ehlers Danlos syndrome | X |  |  |
| Mild persistent asthma | X |  |  |
| Numbness and tingling hands | X |  |  |
| Rib lesion | X |  |  |
| Frequent fevers |  | X |  |

**Table S2.**
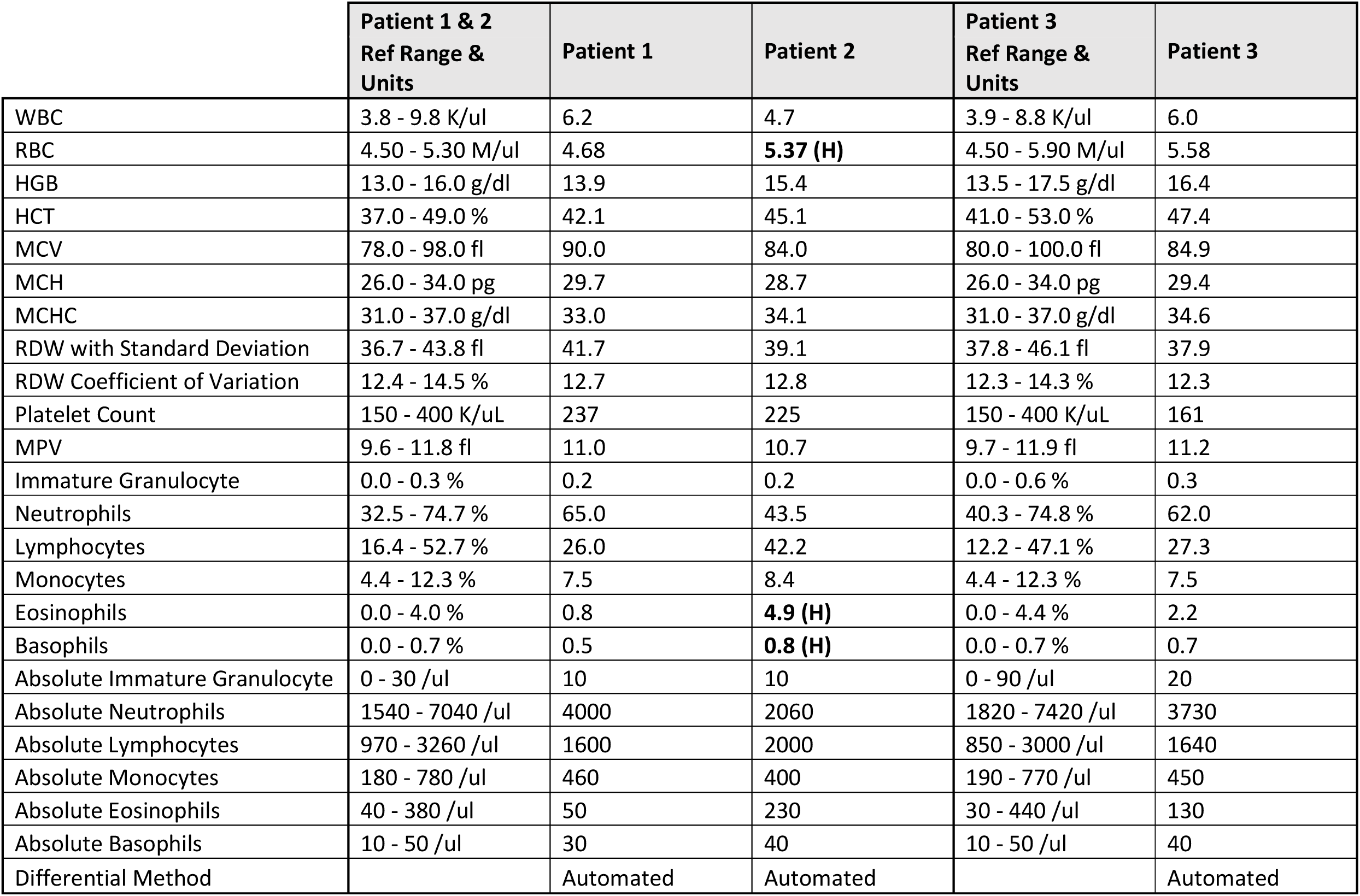
Complete blood count with differential of siblings.

|  | Patient 1 & 2<br>Ref Range &<br>Units | Patient 1 | Patient 2 | Patient 3<br>Ref Range &<br>Units | Patient 3 |
| --- | --- | --- | --- | --- | --- |
| WBC | 3.8 - 9.8 K/ul | 6.2 | 4.7 | 3.9 - 8.8 K/ul | 6.0 |
| RBC | 4.50 - 5.30 M/ul | 4.68 | **5.37 (H)** | 4.50 - 5.90 M/ul | 5.58 |
| HGB | 13.0 - 16.0 g/dl | 13.9 | 15.4 | 13.5 - 17.5 g/dl | 16.4 |
| HCT | 37.0 - 49.0 % | 42.1 | 45.1 | 41.0 - 53.0 % | 47.4 |
| MCV | 78.0 - 98.0 fl | 90.0 | 84.0 | 80.0 - 100.0 fl | 84.9 |
| MCH | 26.0 - 34.0 pg | 29.7 | 28.7 | 26.0 - 34.0 pg | 29.4 |
| MCHC | 31.0 - 37.0 g/dl | 33.0 | 34.1 | 31.0 - 37.0 g/dl | 34.6 |
| RDW with Standard Deviation | 36.7 - 43.8 fl | 41.7 | 39.1 | 37.8 - 46.1 fl | 37.9 |
| RDW Coefficient of Variation | 12.4 - 14.5 % | 12.7 | 12.8 | 12.3 - 14.3 % | 12.3 |
| Platelet Count | 150 - 400 K/uL | 237 | 225 | 150 - 400 K/uL | 161 |
| MPV | 9.6 - 11.8 fl | 11.0 | 10.7 | 9.7 - 11.9 fl | 11.2 |
| Immature Granulocyte | 0.0 - 0.3 % | 0.2 | 0.2 | 0.0 - 0.6 % | 0.3 |
| Neutrophils | 32.5 - 74.7 % | 65.0 | 43.5 | 40.3 - 74.8 % | 62.0 |
| Lymphocytes | 16.4 - 52.7 % | 26.0 | 42.2 | 12.2 - 47.1 % | 27.3 |
| Monocytes | 4.4 - 12.3 % | 7.5 | 8.4 | 4.4 - 12.3 % | 7.5 |
| Eosinophils | 0.0 - 4.0 % | 0.8 | **4.9 (H)** | 0.0 - 4.4 % | 2.2 |
| Basophils | 0.0 - 0.7 % | 0.5 | **0.8 (H)** | 0.0 - 0.7 % | 0.7 |
| Absolute Immature Granulocyte | 0 - 30 /ul | 10 | 10 | 0 - 90 /ul | 20 |
| Absolute Neutrophils | 1540 - 7040 /ul | 4000 | 2060 | 1820 - 7420 /ul | 3730 |
| Absolute Lymphocytes | 970 - 3260 /ul | 1600 | 2000 | 850 - 3000 /ul | 1640 |
| Absolute Monocytes | 180 - 780 /ul | 460 | 400 | 190 - 770 /ul | 450 |
| Absolute Eosinophils | 40 - 380 /ul | 50 | 230 | 30 - 440 /ul | 130 |
| Absolute Basophils | 10 - 50 /ul | 30 | 40 | 10 - 50 /ul | 40 |
| Differential Method |  | Automated | Automated |  | Automated |

**Table S3.** B-T NK panel of siblings.

|  | Patient 1<br>Ref Range & Units | Patient<br>1 | Patient 2<br>Ref Range & Units | Patient<br>2 | Patient 3<br>Ref Range & Units | Patient<br>3 |
| --- | --- | --- | --- | --- | --- | --- |
| CD3+ | 55 - 83 % | 71.0 | 52 - 78 % | 67.8 | 55 - 83 % | 67.3 |
| CD3+ Absolute | 700 - 2100 cells/μl | 1136 | 800 - 3500 cells/μl | 1356 | 700 - 2100 cells/μl | 1104 |
| CD3+/CD4+ | 28 - 57 % | 28.2 | 25 - 48 % | 33.8 | 28 - 57 % | 34.7 |
| CD3+/CD4+ Absolute | 300 -1400 cells/μl | 451 | 400 - 2100 cells/μl | 676 | 300 -1400 cells/μl | 569 |
| CD3+/CD8+ | 10 - 39 % | 32.6 | 9 - 35 % | 29.1 | 10 - 39 % | 26.4 |
| CD3+/CD8+ Absolute | 200 - 900 cells/μl | 522 | 200 - 1200 cells/μl | 582 | 200 - 900 cells/μl | 433 |
| CD19+ | 6 - 19 % | 15.7 | 8 - 24 % | 14.5 | 6 - 19 % | 13.3 |
| CD19+ Absolute | 100 - 500 cells/μl | 251 | 200 - 3500 cells/μl | 290 | 100 - 500 cells/μl | 218 |
| CD20+ | 3.1 - 21.5 % | 15.5 | 3.1 -21.5 % | 14.3 | 3.1 - 21.5 % | 13.2 |
| CD20+ Absolute | 0 - 479 cells/μl | 248 | 0 - 479 cells/μl | 286 | 0 - 479 cells/μl | 216 |
| CD3-/CD16(B73.1)+ and/or CD56+ | 7 - 31 % | 11.9 | 6 - 27 % | 17.3 | 7 - 31 % | 19.3 |
| CD3-/CD16(B73.1)+ and/or CD56+ Abs | 90 - 600 cells/μl | 190 | 70 - 1200 cells/μl | 346 | 90 - 600 cells/μl | 317 |
| CD4+/CD45RA+ | 12.5 - 42.2 % | **9.8 (L)** | 12.5 - 42.2 % | 13.0 | 12.5 - 42.2 % | **11.7 (L)** |
| CD4+/CD45RA+ Absolute | 41 - 1121 cells/μl | 157 | 41 - 1121 cells/μl | 260 | 41 - 1121 cells/μl | 192 |
| CD4+/CD45R0+ | 10.1 - 27.9 % | 18.2 | 10.1 - 27.9 % | 19.2 | 10.1 - 27.9 % | 22.6 |
| CD4+/CD45R0+ Absolute | 153 - 582 cells/μl | 291 | 153 - 582 cells/μl | 384 | 153 - 582 cells/μl | 371 |
| CD4+/CD8+ Ratio | 1 - 3.6 | **0.9 (L)** | 0.9 - 3.4 | 1.2 | 1 - 3.6 | 1.3 |

**Table S4.** Clinical test of lymphocyte proliferation in response to antigens/mitogens.

|  | Patient 1 |  | Patient 2 |  | Patient 3 |  |
| --- | --- | --- | --- | --- | --- | --- |
|  | CPM | SI | CPM | SI | CPM | SI |
| Antigen background | 86 |  | 227 |  | 4745 |  |
| Tetanus | 5070 | 59 | 9215 | 41 | 32298 | 7 |
| Candida | 3616 | 42 | 6056 | 27 | 14139 | 3 |
| Mitogen background | 172 |  | 266 |  | 395 |  |
| PWM | 20233 | 118 | 20210 | 76 | 81204 | 206 |
| ConA | 54679 | 318 | 53397 | 201 | 100903 | 255 |
| PHA | 62156 | 361 | 64221 | 241 | 171802 | 435 |
|  | Control of Patient 1 |  | Control of Patient 2 |  | Control of Patient 3 |  |
|  | CPM | SI | CPM | SI | CPM | SI |
| Antigen background | 553 |  | 553 |  | 799 |  |
| Tetanus | 15309 | 28 | 15309 | 28 | 27475 | 34 |
| Candida | 12932 | 23 | 12932 | 23 | 28674 | 36 |
| Mitogen background | 299 |  | 299 |  | 556 |  |
| PWM | 51001 | 171 | 51001 | 171 | 72116 | 130 |
| ConA | 60878 | 204 | 60878 | 204 | 130355 | 234 |
| PHA | 77314 | 259 | 77314 | 259 | 186396 | 336 |

**Table S5.** EBV viral loads and EBV antibodies in two of the three siblings.

|  | Patient 2 | Patient 3 | Negative reference range |
| --- | --- | --- | --- |
| **EBV antibody detection** | U/ml | U/ml | U/ml |
| early antigen IgG | 18.1 | 14.7 | 0.0 - 8.9 |
| nuclear antigen IgG | 160 | 339 | 0.0 - 17.9 |
| VCA IgG | 85.1 | 519 | 0.0 - 17.9 |
| VCA IgM | <36.0 | <36.0 | 0.0 - 35.9 |
| **EBV quantitative PCR** | - | - | - |
| viral load, blood | 350 | n/a | <125 |
| log value, blood | 2.54 | n/a | <2.1 |
| viral load, lymphs | n/a | 83 | <50 |
| viral load, plasma | n/a | 581 | <50 |
| log value, lymphs | n/a | 1.92 | <1.7 |
| log value, plasma | n/a | 2.76 | <1.7 |

**Table S6.** Summary of known genetic variants in the patients and their mother.

|  |  |  | Patient 1 | Patient 2 | Patient 3 | Mother |
| --- | --- | --- | --- | --- | --- | --- |
| Gene | Variant | dbSNP |  |  |  |  |
| **Pathogenic variants** |  |  |  |  |  |  |
| CACNA1F | p.R625X | rs886039559 | - / 0 | - / 0 | - / 0 | - / + |
| **Benign and likely benign variants** |  |  |  |  |  |  |
| SH2D1A | p.V40M | rs199639961 | + / 0 | - / 0 | + / 0 | - / + |
| AMPD | p.Q45X | rs17602729 | - / - | + / + | - / + | - / + |
| AMPD | p.P81L | rs61752479 | - / - | + / + | - / + | - / + |
| MEFV | p.G138G | rs224224 | + / + | - / + | - / + | N / A |
| MEFV | p.A165A | rs224223 | + / + | - / + | - / + | N / A |
| MEFV | p.D102D | rs224225 | + / + | - / + | - / + | N / A |
| MEFV | p.R314R | rs224213 | - / - | - / + | - / + | N / A |
| MEFV | p.E474E | rs224208 | - / - | - / + | - / + | N / A |
| MEFV | p.Q476Q | rs224207 | - / - | - / + | - / + | N / A |
| MEFV | p.D510D | rs224206 | - / - | - / + | - / + | N / A |
| MEFV | p.P588P | rs1231122 | - / - | - / + | - / + | N / A |
| NLRP3 | p.A244A | rs3806268 | - / - | - / - | - / - | N / A |
| TNFRSF1A | p.P12P | rs767455 | - / - | - / + | - / + | N / A |
| TNFRSF1A | c.625+10 | rs1800693 | - / - | - / + | - / + | N / A |
| TNFRSF1A | c.740-9 | rs12426675 | + / + | - / + | - / + | N / A |
| **Genes reviewed for secondary findings** |  |  |  |  |  |  |
| ACTA2 |  |  | + / + | + / + | + / + | N / A |
| ACTC1 |  |  | + / + | + / + | + / + | N / A |
| APC |  |  | + / + | + / + | + / + | N / A |
| APOB |  |  | + / + | + / + | + / + | N / A |
| BRCA1 |  |  | + / + | + / + | + / + | N / A |
| BRCA2 |  |  | + / + | + / + | + / + | N / A |
| CACNA1S |  |  | + / + | + / + | + / + | N / A |
| COL3A1 |  |  | + / + | + / + | + / + | N / A |
| DSC2 |  |  | + / + | + / + | + / + | N / A |
| DSG2 |  |  | + / + | + / + | + / + | N / A |
| DSP |  |  | + / + | + / + | + / + | N / A |
| FBN1 |  |  | + / + | + / + | + / + | N / A |
| GLA |  |  | + / + | + / + | + / + | N / A |
| KCNH2 |  |  | + / + | + / + | + / + | N / A |
| KCNQ1 |  |  | + / + | + / + | + / + | N / A |
| LDLR |  |  | + / + | + / + | + / + | N / A |
| LMNA |  |  | + / + | + / + | + / + | N / A |
| MEN1 |  |  | + / + | + / + | + / + | N / A |
| MLH1 |  |  | + / + | + / + | + / + | N / A |
| MSH2 |  |  | + / + | + / + | + / + | N / A |
| MSH6 |  |  | + / + | + / + | + / + | N / A |
| MUTYH |  |  | +/+ | +/+ | +/+ | N/A |
| MYBPC3 |  |  | +/+ | +/+ | +/+ | N/A |
| MYL2 |  |  | +/+ | +/+ | +/+ | N/A |
| MYL3 |  |  | +/+ | +/+ | +/+ | N/A |
| MYLK |  |  | +/+ | +/+ | +/+ | N/A |
| MYH7 |  |  | +/+ | +/+ | +/+ | N/A |
| MYH11 |  |  | +/+ | +/+ | +/+ | N/A |
| NF2 |  |  | +/+ | +/+ | +/+ | N/A |
| PCSK9 |  |  | +/+ | +/+ | +/+ | N/A |
| PKP2 |  |  | +/+ | +/+ | +/+ | N/A |
| PMS2 |  |  | +/+ | +/+ | +/+ | N/A |
| PRKAG2 |  |  | +/+ | +/+ | +/+ | N/A |
| PTEN |  |  | +/+ | +/+ | +/+ | N/A |
| RB1 |  |  | +/+ | +/+ | +/+ | N/A |
| RET |  |  | +/+ | +/+ | +/+ | N/A |
| RYR1 |  |  | +/+ | +/+ | +/+ | N/A |
| RYR2 |  |  | +/+ | +/+ | +/+ | N/A |
| SCN5A |  |  | +/+ | +/+ | +/+ | N/A |
| SDHAF2 |  |  | +/+ | +/+ | +/+ | N/A |
| SDHB |  |  | +/+ | +/+ | +/+ | N/A |
| SDHC |  |  | +/+ | +/+ | +/+ | N/A |
| SDHD |  |  | +/+ | +/+ | +/+ | N/A |
| SMAD3 |  |  | +/+ | +/+ | +/+ | N/A |
| STK11 |  |  | +/+ | +/+ | +/+ | N/A |
| TGFBR1 |  |  | +/+ | +/+ | +/+ | N/A |
| TGFBR2 |  |  | +/+ | +/+ | +/+ | N/A |
| TMEM43 |  |  | +/+ | +/+ | +/+ | N/A |
| TNNI3 |  |  | +/+ | +/+ | +/+ | N/A |
| TNNT2 |  |  | +/+ | +/+ | +/+ | N/A |
| TP53 |  |  | +/+ | +/+ | +/+ | N/A |
| TPM1 |  |  | +/+ | +/+ | +/+ | N/A |
| TSC1 |  |  | +/+ | +/+ | +/+ | N/A |
| TSC2 |  |  | +/+ | +/+ | +/+ | N/A |
| VHL |  |  | +/+ | +/+ | +/+ | N/A |
| WT1 |  |  | +/+ | +/+ | +/+ | N/A |

+/+ normal homozygous, -/- mutant homozygous, -/+ mutant heterozygous,
+ / 0 normal hemizygous, - / 0 mutant hemizygous

**Table S7.**
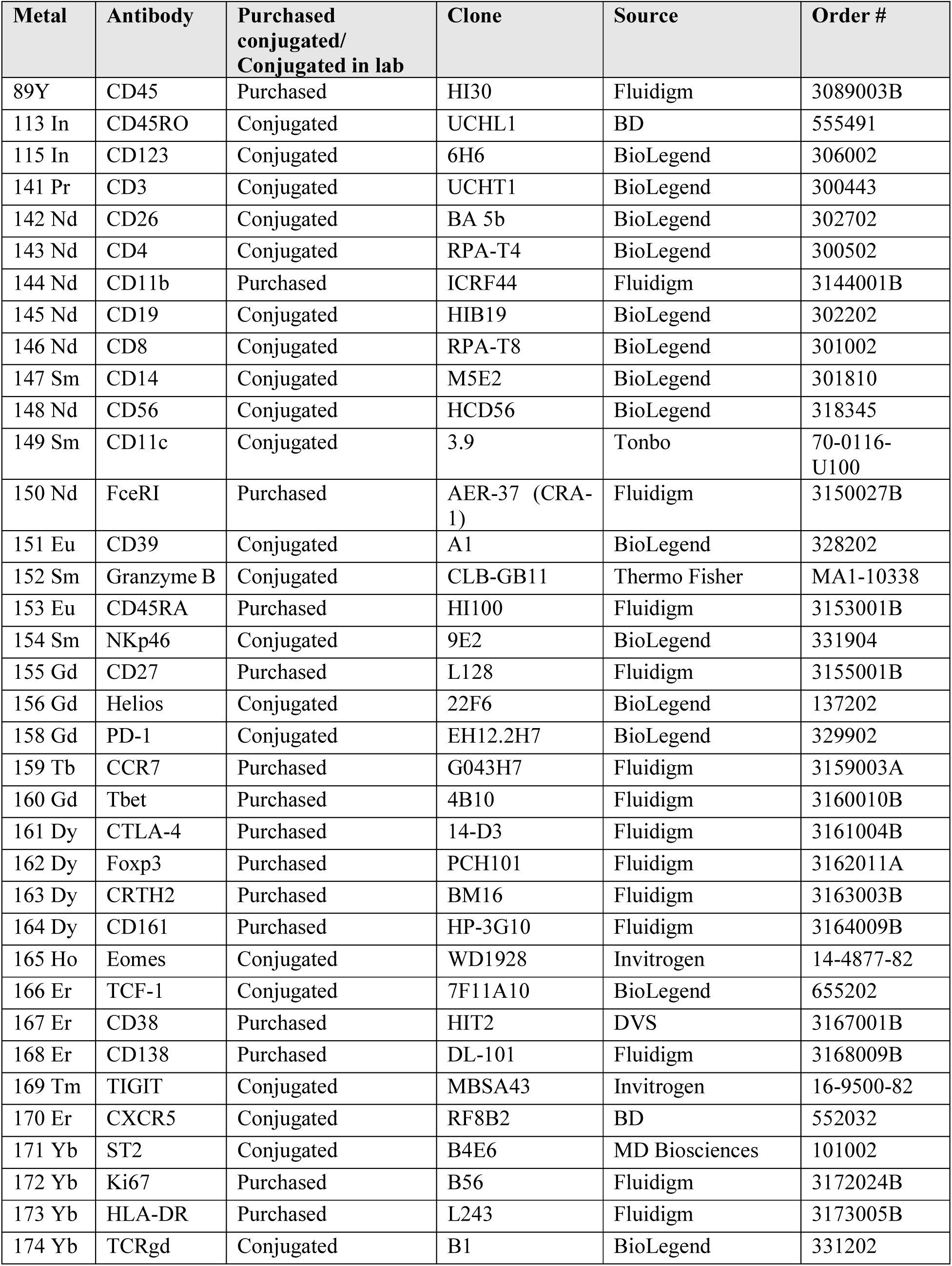
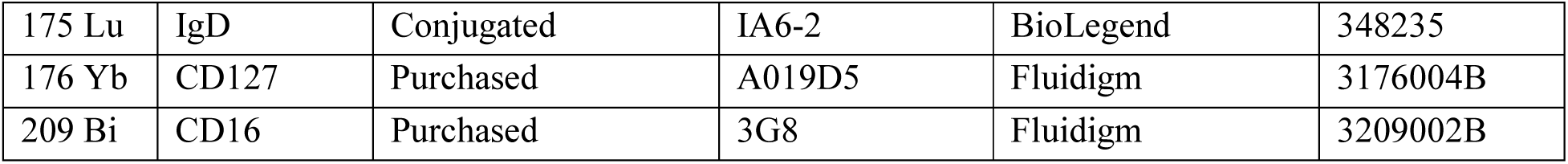
Mass cytometry (CyTOF) human panel.

**Table S8.** Flow cytometry human cytokine panel.

| Fluorochrome | Antibody | Clone | Source | Order # |
| --- | --- | --- | --- | --- |
| BV421 | IL-2 | 5344.111 | BD | 562914 |
| V500 | CD14 | M5E2 | BD | 561391 |
| V500 | CD16 | 3G8 | BD | 561394 |
| V500 | CD19 | HIB19 | BD | 561121 |
| Aqua | Live/Dead |  | Invitrogen | L34966 |
| QD585 | CD3 | OKT3 | BioLegend | 317304 |
| BV605 | CD45RA | HI100 | BioLegend | 304134 |
| SB645 | CD8 | RPA-T8 | Invitrogen | 64-0088-42 |
| BV711 | CD279/PD-1 | EH12.1 | BD | 564017 |
| BV785 | CD27 | O323 | BioLegend | 302831 |
| AF647 | IL-21 | 3A3-N2 | BioLegend | 513006 |
| AF700 | Foxp3 | PCH101 | Invitrogen | 56-4776-41 |
| APC/Cy7 | IL-17A | BL168 | BioLegend | 512319 |
| AF488 | IL-10 | JES3-9D7 | BioLegend | 501411 |
| PE | IL-13 | JES10-5A2 | BioLegend | 501903 |
| PE-TxRd | CD38 | HIT2 | Life | MHCD3817 |
| PE-Cy5.5 | CD4 | SK3 | eBio | 35-0047-42 |
| PE/Cy7 | IFNg | B27 | BioLegend | 506518 |

